# Feature extraction approach in single-cell gene expression profiling for cell-type marker identification

**DOI:** 10.1101/686659

**Authors:** Nigatu A. Adossa, Leif Schauser, Vivi G. Gregersen, Laura L. Elo

## Abstract

**Background:** Recent advances in single-cell gene expression profiling technology have revolutionized the understanding of molecular processes underlying developmental cell and tissue differentiation, enabling the discovery of novel cell-types and molecular markers that characterize developmental trajectories. Common approaches for identifying marker genes are based on pairwise statistical testing for differential gene expression between cell-types in heterogeneous cell populations, which is challenging due to unequal sample sizes and variance between groups resulting in little statistical power and inflated type I errors.

**Results:** We developed an alternative feature extraction method, Marker gene Identification for *Cell-type I*dentity (***MICTI***) that encodes the cell-type specific expression information to each gene in every single-cell. This approach identifies features (genes) that are cell-type specific for a given cell-type in heterogeneous cell population. To validate this approach, we used (i) simulated single cell RNA-seq data, (ii) human pancreatic islet single-cell RNA-seq data and (iii) a simulated mixture of human single-cell RNA-seq data related to immune cells, particularly B cells, CD4+ memory cells, CD8+ memory cells, dendritic cells, fibroblast cells, and lymphoblast cells. For all cases, we were able to identify established cell-type-specific markers.

**Conclusions:** Our approach represents a highly efficient and fast method as an alternative to differential expression analysis for molecular marker identification in heterogeneous single-cell RNA-seq data.

## Background

### Single-cell RNA-seq protocols

Technical advances in single-cell RNA-sequencing have enabled RNA-seq as the method of choice for studying organ and cellular differentiation processes and cell heterogeneity in complex mixtures found in body fluids and tissues. SMART-seq2 (Picelli et al., 2014), CELL-seq (Hashimshony, Wagner, Sher, & Yanai, 2012) and Drop-seq (Klein & Macosko, 2017) are among widely used protocols for single-cell expression profiling. In general, single cells are obtained using Fluorescence-activated cell sorting (FACS) (Nybo, 2010) or microfluidics-based techniques (Klein & Macosko, 2017). Once individual cells are isolated, the RNA-seq protocols allow the quantification of gene expression. Single-cell protocols often involve unique molecular identifiers (UMIs), which uniquely tag a captured messenger RNA transcript. This greatly improves the accuracy of expression measurement by removing the amplification biases. Although this technology is powerful in studying the stochasticity of gene expression in development and identifying novel cell-types, the data analysis challenges remain.

### Noise in Single-cell RNA-seq

As the number of transcripts coming from a single cell is scanty compared to bulk RNA extraction, single-cell RNA-seq data is more prone to amplification biases and sampling effects during transcription capture. Therefore, it is very important to control the quality of the single-cell data prior to embarking on downstream analysis. Quality control can be done on the number of reads in each cell. Read quality control (QC) software such as FASTQC (Andrews, 2010) and RSeQC (X. Li, Nair, Wang, & Wang, 2015) are used in bulk as well as single-cell RNA-seq data. Besides quality control at raw read level, it is also important to control quality in terms of transcripts captured per cell. Cells with low gene capture rates should be removed as failure to do so can heavily affect the downstream analysis. SinQC (Jiang, Thomson, & Stewart, 2016) is a quality control software specifically designed for single-cell RNA-seq data by integrating both data quality information and gene expression patterns. Another source of noise is the inclusion of cells undergoing cell apoptosis induced by the single-cell harvesting protocol. A criterion that has been successfully used as the measure of cell viability is the proportion of reads mapping to the mitochondrial genes. If most of the reads are mapped to mitochondrial genes, this indicates the leaking or degradation of cytoplasmic RNA in apoptotic cells (Bacher & Kendziorski, 2016). Hence, many protocols use the proportion of reads mapped to artificially spiked-in RNAs as a robust workflow quality control measure (Chen et al., 2016).

### Normalization in single-cell RNA-seq data

Due to the low number of transcripts available in a single cell and experimental limits to transcript sampling rates, single-cell RNA-seq data are zero-inflated and sparse (Vu et al., 2016). Normalization is performed to account for systematic biases in capture rates related to technical and undesired biological effects. Bayesian probabilistic models of expression-magnitude distortions of typical single-cell RNA sequencing data consider high-level noise and intrinsic biological variation (Kharchenko, Silberstein, & Scadden, 2014). A similar method also considers sampling noise and global cell-to-cell variation in sequencing efficiency as “noise models” in single-cell RNA-seq data (Grün, Kester, & Van Oudenaarden, 2014). A two-part generalized linear model to parameterize the bimodal nature of single cell measurements was developed by (Finak et al., 2015). (Buettner et al., 2015) has used a latent variable model to account for the sources of variation in single-cell gene expression data.

However, normalization methods developed for bulk RNA-seq data are also used for single-cell RNA-seq normalization even though some researchers argue that such an approach is error-prone and heavily influences the downstream analysis (Vallejos, Risso, Scialdone, Dudoit, & Marioni, 2017). For example, DEseq2 is a model based differential expression analysis method designed for the bulk RNA-seq data using negative binomial (NB) distribution. Normalization in DESeq2 is performed by estimating the global scaling factor to normalize the given raw count data (Kvam, Liu, & Si, 2012). Transcripts per million reads (TPM) and reads per kilobase per millions of mapped read (RPKM) provide per-single-cell RNA-seq normalization measures. The trimmed mean of M-values normalization (TMM) (Robinson & Oshlack, 2010) is a scaling factor based method normalizing across all single-cells in a given experiment. These approaches can bias the downstream analysis through the effect of a few highly expressed genes. To avoid the pitfalls, normalization based on artificial spike-in molecules in single-cell RNA-seq experiments has been developed. Efficient and robust methods single-cell RNA-seq normalization method are currently under development (Vallejos et al., 2017).

### Cell-type identification

The main use of single-cell RNA-sequencing is to study cellular heterogeneity in tissues or organs and identify events that govern cell development and differentiation. Unsupervised clustering methods have been used for the identification of cell-types from normalized single-cell RNA-seq data. SC3 (Kiselev et al., 2016) offers several clustering algorithms and brings consensus clusters to identify cell-types. Seurat (Satija, Farrell, Gennert, Schier, & Regev, 2015) uses graph-based clustering (Levine et al., 2015) methods together with modularity optimization techniques (Blondel, Guillaume, Lambiotte, & Lefebvre, 2008) for optimization of clusters. Monocle (Trapnell et al., 2014) and TSCAN (Ji & Ji, 2016) extract features of the gene expression profile of each cell by independent component analysis (ICA) followed by a minimum spanning tree (MST) algorithm to visualize the cell-types as a developmental trajectory in pseudo-time. Density (Haghverdi, Buettner, & Theis, 2015) uses diffusion map for visualization of cellular trajectory in development.

### Marker genes deriving cell identity

Once cell-types are identified, the next challenge is to pinpoint the source of heterogeneity in cells. Markers are genes with altered expression level as a function cell-type or genes that change in expression level as a function of development or response to a disease state. To identify such markers, single-cell RNA-seq analysis pipelines such as MAST (Finak et al., 2015) and BPSC (Vu et al., 2016) use differential gene expression analysis (DGE) methods developed specifically for single-cell RNA-seq data using two-part generalized linear model and beta-Poisson mixture model respectively. Many single-cell RNA-seq analysis pipelines make use of methods that were developed for bulk RNA-seq data such as DESeq2 (Love, Anders, & Huber, 2014). ROTS (Suomi, Seyednasrollah, Jaakkola, Faux, & Elo, 2017), a data distribution independent statistical testing method, is another pipeline designed to identify DEGs by adjusting the test statistics according to the nature of the dataset.

In a single-cell RNA-seq experiment, the number of cells per cell-type is usually variable. This leads to reduced statistical power when attempting to identify differentially expressed marker genes. Comparisons of various differential expression analysis methods for single-cell RNA-seq data have shown that they yield different results both in terms of the number of detected DEG and consistency of DEG’s identity across the methods (Miao & Zhang, 2016). Performing pairwise differential expression analysis to identify cell-type marker genes in heterogeneous single-cell RNA-seq data is inefficient in terms of execution time and redundant use, especially when considering several cell-types in the heterogeneous cell population.

Alternative approaches to the problem make use of feature extraction, where information extraction methods are used to mine higher-level information from the datasets resulting in non-redundant information that can be used for high accuracy cell-type identification. Feature extraction is widely used for dimensionality reduction such as principal component analysis (PCA) (Breu, Guggenbichler, & Wollmann, 2008), independent component analysis (ICA) (Kachenoura, Albera, Senhadji, & Comon, 2008), non-negative matrix factorization (NMF) (Eggert & Krner, 2004), linear discriminate analysis (LDA) (M. Li & Yuan, 2005) and autoencoders (Wang, Huang, Wang, & Wang, 2014). For document classification and sentiment analysis, term frequency-inverse document frequency (TF-IDF) (Ramos, 2003) and latent Dirichlet allocation (LDA) (Blei et al., 2003) are widely used feature extraction methods.

Here we adopt the TF-IDF method to develop ***MICTI*** (Fig. 1), a feature extraction method for Marker gene Identification for *Cell-type* Identity in single-cell RNA-seq data. ***MICTI*** can be used as an alternative method with differential gene expression for cell-type specific marker identification in single-cell RNA-seq data. The aim of ***MICTI*** is to rapidly identify key cell-type (or cluster) specific genes for the analysis of heterogeneous single-cell RNA-seq. Making use of the sparsity of the normalized expression data, the cell-type (or cluster) specific information is encoded for each gene in every cell. This allows ***MICTI*** to avoid variance based gene filtering steps typically used in the preprocessing of single-cell RNA-seq analysis.

**Figure 1.**
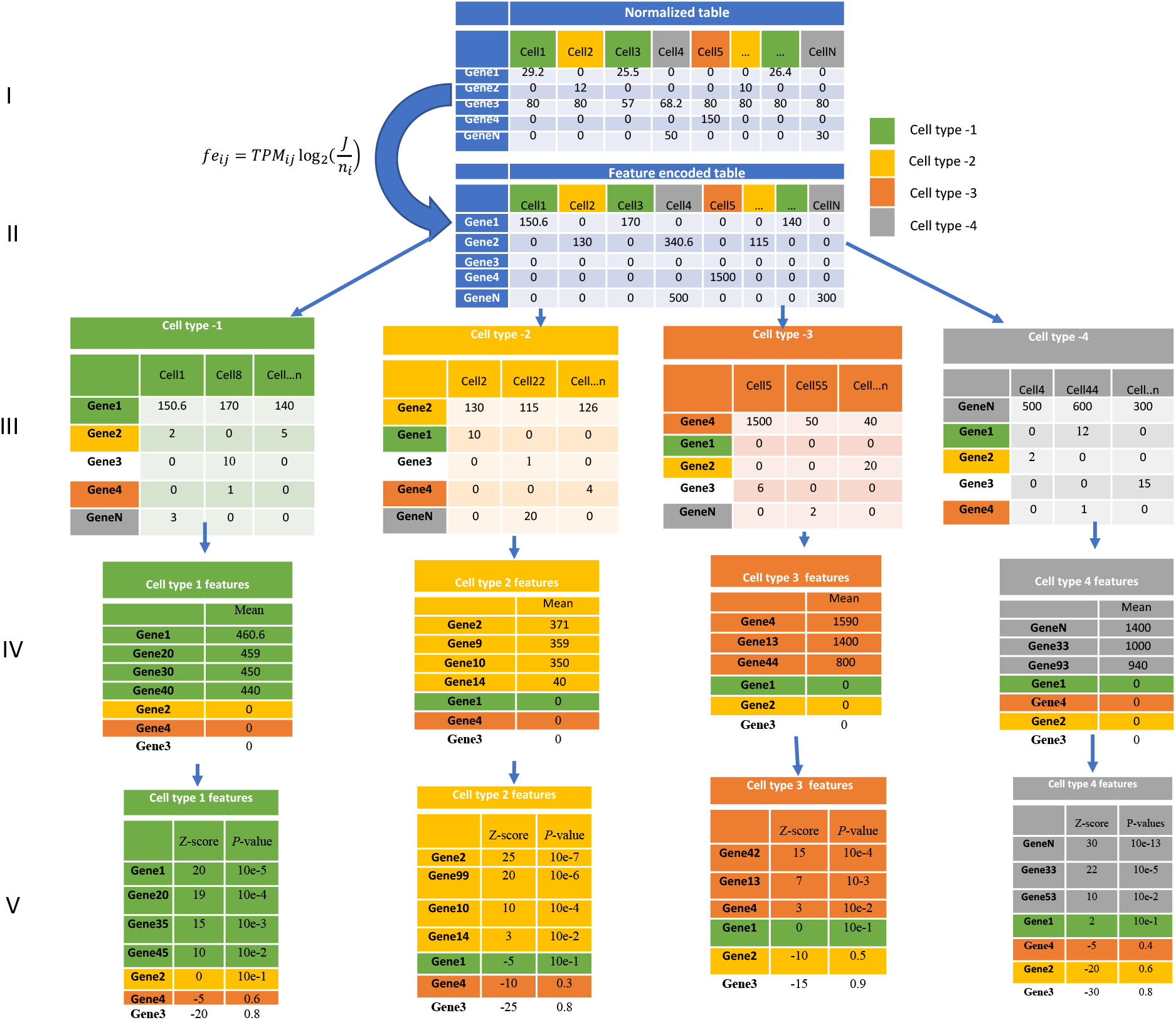
Workflow for *MICTI*. **(I)** raw expression matrixes are normalized to account for differences in library size. (II) Then expression values are transformed into feature (cell-type) specific values by feature encoding for each of the gene in every cell. **(III)** The cells are clustered by cell-types. **(IV)** The cluster mean feature encoded expression values for each gene is normalized by expression variance within the cluster in a logarithmic scale. **(V)** Finally, the p-value is calculated for cell-type (or cluster) marker genes.

The ***MICTI*** workflow comprises of five analytical steps (Fig. 1). The first step is pre-processing and normalization (Fig. 1I). Secondly, the normalized expression data is transformed into cell-type specific feature-encoded expression data by using scarcity of expression factor (Fig. 1II). In the third step, cells are grouped or clustered based on their cell-types (Fig. 1III). Then in the fourth step (Fig. 1IV), the cluster mean feature-encoded expression is calculated for each of the genes. Finally, the p-value is calculated for each of the genes (Fig. 1V) in each of the clusters normalizing the cluster mean values with their within-cluster variance. The genes with the highest average expression and lowest within-cluster variance hit the significance threshold identifying it as a marker.

We demonstrate the efficiency of ***MICTI*** in three case studies. First, we analysed a simulated single-cell RNA-seq dataset (*supplementary file 1*). Second, we analysed a publicly available human pancreatic islets data from Gene Expression Omnibus (*supplementary file 2*). Third, we analysed an artificially created mixture of single-cell RNA-seq data from six immune cell expression data from Gene Expression Omnibus, where we selected 1153 cells sampled from different tissues with different disease conditions for CD4+ memory cells, CD8+ memory cells, B cells, Dendritic cells, Fibroblasts and Lymphoblast cells (*Table 1*).

**Table 1.** Single-cell RNA-seq datasets from GEO that were used for creating the artificial mixture of single-cell data.

| Project Id | Selected cells | Tissue |
| --- | --- | --- |
| GSE75748(1810 cells) | Fibroblast(159 cells) | Foreskin |
| GSE81861(1220 cells) | Lymphoblast(59 cells) | Bone marrow |
| GSE44618(62 cells),<br>GSE81861(1220 cells) | B cell(174 cells) | Peripheral blood |
| GSE96562(149 cells),<br>GSE96563(9 cells),<br>GSE96568(246 cells) | CD4+ memory T cell(393 cells) | Peripheral blood |
| GSE85527(219 cells),<br>GSE85530(63 cells),<br>GSE96564(45 cells) | CD8+ memory T cell (263 cells) | Peripheral blood |
| GSE89232(957 cells) | Conventional dendritic cell(105 cells) | Peripheral blood |

## Results

### Case study 1: Simulated single cell RNA-seq data

We used SymSim (Zhang, Xu, & Yosef, 2019), a software that explicitly models the processes that give rise to data observed in single-cell RNA-seq experiments, for simulating 200 cells with five distinct single cell populations. The parameters used in the simulation process resemble the actual single-cell RNA-seq data generation process. The input parameters include, for example, mRNA capture rate, the number of PCR cycles, sequencing depth, or the use of unique molecular identifiers.

Fig. 2b shows the 2D visualization for the simulated single-cell RNA-seq data. The hierarchical similarity of the cell-types in the simulated dataset is in accordance with the input tree parameter (Fig. 2a). Cell-types A and B are clustered together and cell-types C and D are clustered together. The cell-type E is relatively distinct as compared to the other cell-types. Fig. 2c shows a heatmap for the markers identified by ***MICTI*** with *p-value* less than 0.01 and Z-score greater than 0. The cell-type 1 marker genes distinctly have high expression in cell-type 1 cells and low expression in the other cell-types. Markers for cell-type A and B share common marker genes. The same is true for cell-type C and D cells. This bolsters the hypothesis that similar cell-types have similar gene expression pattern and marker genes.

**Figure 2.**
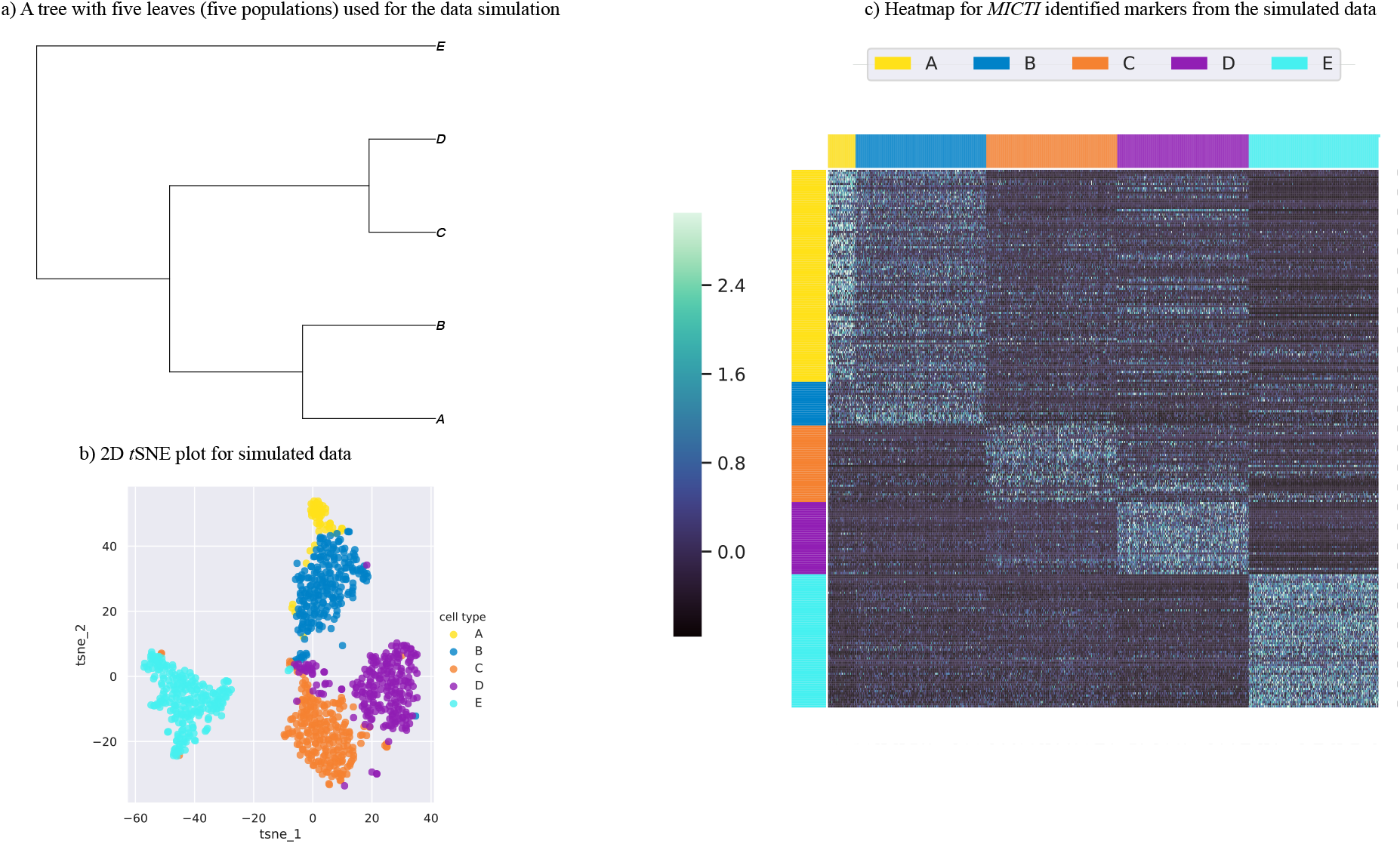
*MICTI* marker gene identification on simulated single-cell RNA-seq data. (**a**) A tree with five leaves or populations (A to E) used as an input for the data simulation. (**b**) 2D t-SNE plot for the simulated single-cell RNA-seq data. (**c**) Heatmap for the ***MICTI*** identified markers from the simulated data. The Column represent the cells with cell-types defined by the colour code in the legend and the row is the expression of marker genes identified by ***MICTI***. The row colour coding given in the legend represents the cell-type specific marker genes.

### Case study 2: Human pancreatic islet cell-type markers

We next applied *MICTI* on real single-cell RNA-seq dataset. We used publicly available human pancreatic islet single-cell RNA-seq data with the GEO accession number of GSE86469. Pancreatic islets are cells that play a crucial role in controlling the blood glucose level (Lawlor et al., 2017). Human islets consist mostly of beta, alpha, delta, stellate, ductal, acinar and gamma/pancreatic polypeptide (PP) cells (Lawlor et al., 2017). Changes in the proportion and function of these cell-types are associated with the genetic and pathophysiology of monogenic type 1 and type 2 diabetics (Lawlor et al., 2017).

We analysed 638 pancreatic islet cells (*supplement file 2*) with 26,616 genes. TPM normalization was used before the ***MICTI*** marker gene identification analysis. Labelling of cells into cell-types was performed using the metadata information associated with the dataset and Fig. 3a shows the lower dimensional representation using t-SNE plot. The heatmap for the expression of the cell-type marker genes in each of the pancreatic islet endocrine cell-types is illustrated in Fig. 3b. From the heatmap, it can be shown that cell-type marker genes identified by ***MICTI***, have high expression patterns for their respective clusters or cell-types.

**Figure 3.**
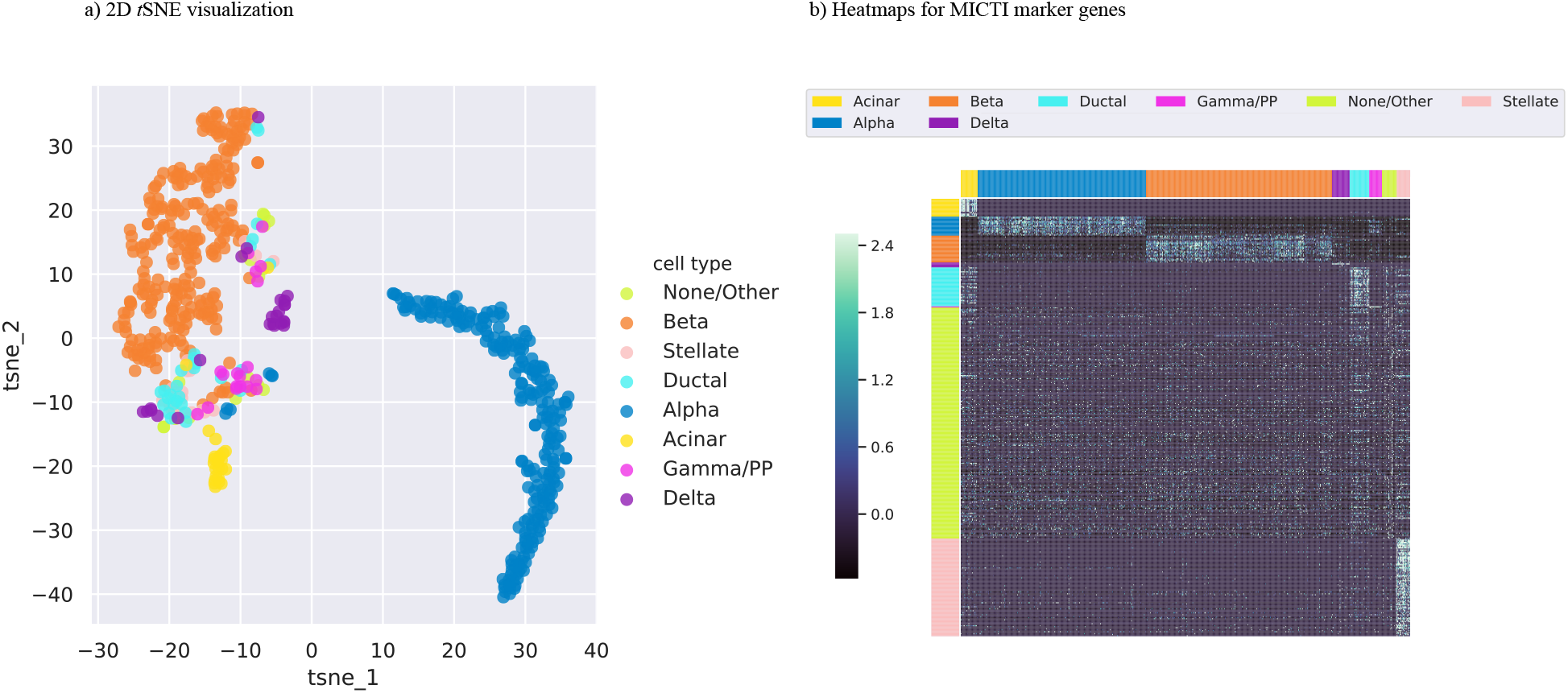
*MICTI* marker gene identification on pancreatic islet single-cell RNA-seq data. *(**a**) 2D visualization of human pancreatic islet single-cell RNA-seq data using* t-SNE *plot. (**b**) The gene expression heatmap for **MICTI*** identified *pancreatic islet cell-type markers. The columns represent the cells sorted by their cell-type and the rows are the marker genes identified by **MICTI**. The colour on the column and row headers corresponds to each of the cell-types as it is illustrated in the legend*.

Genes such as *INS* (beta), *GCG* (alpha), *SST* (delta), *PPY* (PP/gamma), *PRSS1* (acinar), the extracellular matrix protein gene *COL1A1* (stellate) and the structural keratin gene *KRT19* (ductal) were identified as marker genes to their respective cell-types by ***MICTI***. (Lawlor et al., 2017) has also reported all these genes as key cell-type marker genes in their respective cell-types. All identified marker genes with the *p-value* less than 0.01 and Z-score of greater than 0 are provided as a *supplement file 3*.

The gene set enrichment analysis for marker genes of Acinar pancreatic islet (*REG1A, PRSS1, CTRB1, REG1B, CELA3A, CPA1, CTRB2, REG3A, CLPS, PLA2G1B, SYCN, SPINK1*) shows that they are enriched for the KEGG pathway of Pancreatic secretion (KEGG:04972). Pancreatic acinar cells are responsible for the secretion of digestive enzymes (Geron, Schejter, & Shilo, 2014). Alpha pancreatic islet marker genes, for example, *CHGB, PAPPA2, SCG2, SERPINA1*, were enriched for the Reactome pathway term of regulation of Insulin-like Growth Factor (IGF) transport and uptake by Insulin-like Growth Factor Binding Proteins (*IGFBPs*) (*REAC:R-HSA-381426*). The secretory granule is used to release Glucagon by alpha pancreatic islet cells (González-Vélez, Dupont, Gil, González, & Quesada, 2012). The regulation of insulin secretion (*GO:0050796*) was also an enriched GO term for the beta-pancreatic islet cell’s marker genes (*ABCC8, CASR, PFKFB2, RBP4, HADH, G6PC2, SLC30A8, EFNA5*). Insulin processing (REAC:R-HSA-264876) and Insulin secretion (KEGG:04911) were the Reactome and KEEG pathways enriched for the ***MICTI*** identified markers for beta-pancreatic islet cells (*SLC30A8, PCSK1, ABCC8, ADCYAP1, INS*) respectively. *INS* is an insulin producing gene that regulate the glucose level in the blood (Rochet et al., 1997). The ***MICTI*** cell-type specific markers and their gene-list enrichment analysis result is provided in *supplementary file 4*.

### Case study 3: Mixture of public single-cell RNA-seq datasets

Finally, we validated ***MICTI*** using a mixture of heterogeneous public single-cell RNA-seq data generated by different labs. We created an artificial mixture of single cells from 10 human immune single-cell experimental datasets. From these datasets, we selected 1153 cells representing single-cell gene expression measurements for CD4+ memory cells, CD8+ memory cells, B cells, Dendritic cells, Fibroblasts and Lymphoblast (*Table* 1). The TPM normalized expression matrix for 13880 genes in 1153 cells are provided in *supplement file 5*.

The cluster assignment of a cell to its cell-type was based on the metadata information from the corresponding samples. The lower dimensional visualization of cells in 2D *t-SNE* plot is shown in Fig. 4a. All ***MICTI*** marker genes for each of the cell-type are illustrated in *supplementary file 6*. Applying the ***MICTI*** feature extraction approach, we found that genes with the p-value of less than 0.01 were part of pathways or gene ontology terms associated with the given cell-types (*supplementary file 7*).

**Figure 4.**
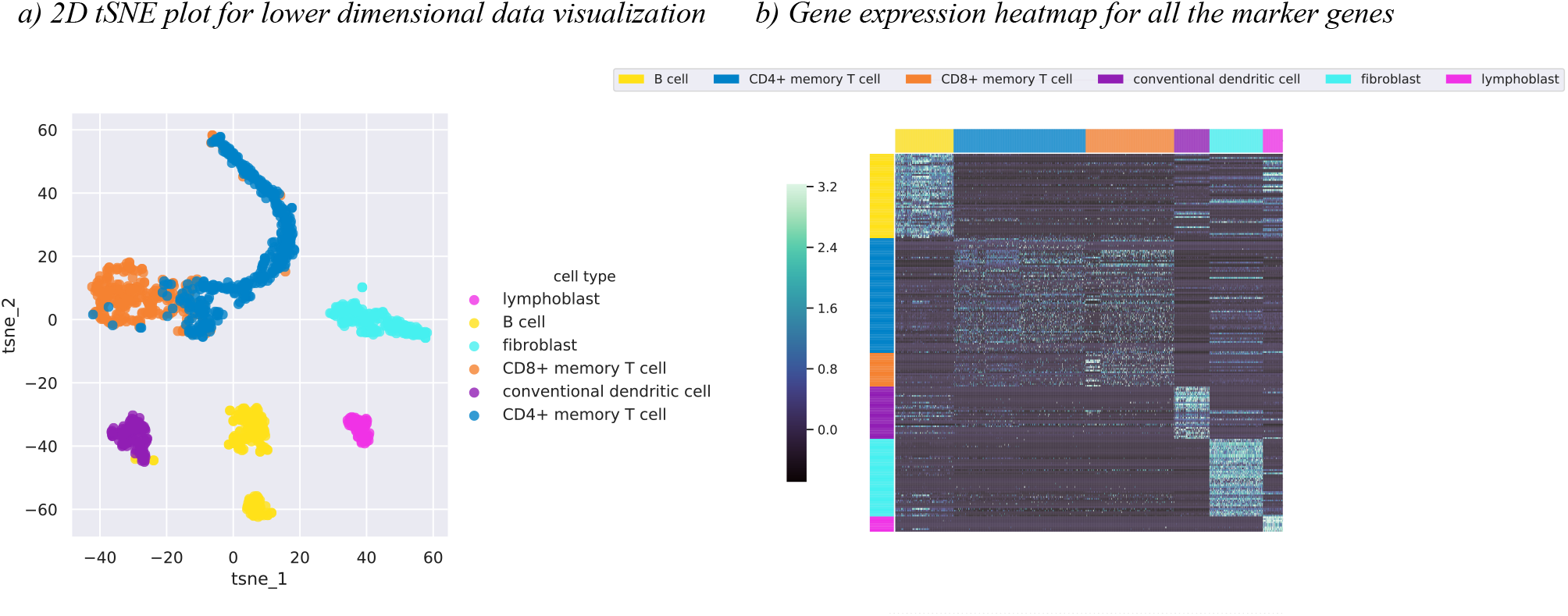
*MICTI* marker gene identification on artificial mixture of single-cell RNA-seq data. (**a**) The 2D visualization of the gene expression dataset from 1153 cells of the six cell-types. (**b**) The gene expression heatmap for MICTI identified marker genes across all cell-types. Columns are the six cell-types as represented in the legend and the rows are the marker genes for each of the cell-types. The row colour coding indicates the cell-type specific marker genes.

The gene expression heatmap (Fig. 4b) for ***MICTI*** identified marker genes show that the expression pattern of the cell-type specific marker genes are specific to the corresponding cell-types. We can also notice that cells that are closely related functionally (B cells and dendritic cells or CD4+ and CD8+ cells) share common cell-type marker genes.

Fig. 5 illustrates the key cell-type specific features (genes) that govern the cell identity for B cells, CD4+ memory cells, CD8+ memory cells, Dendritic cells, Fibroblast cells and Lymphoblast cells. The numbers in the Venn diagram(Fig. 5) indicate the number of marker genes identified in either intersection regions or in the exclusive cell-type regions. Out of the 50 significant cell-type specific genes in the B cells, 31 of them were exclusively B-cell specific genes. The 50 significantly B-cell specific genes were involved in antigen processing and presentation, antigen receptor-mediated signalling pathway, interferon-gamma mediated signalling pathway, cytokine-mediated signalling pathways, cellular response to interferon gamma and MHC class II protein complex binding biological ontology terms among others (*supplement file 7*). These pathways and ontology terms are B cell related. (X. Chen & Jensen, 2008) has shown that B cells are professional antigen-presenting cells(APCs). 10 genes were common cell-type specific genes that were expressed in both B cells and Dendritic cells. These genes were also enriched for antigen processing and presentation. It is known that both B cells and Dendritic cells are antigen-presenting cells (APCs) (Segura & Villadangos, 2009).

**Figure 5.**
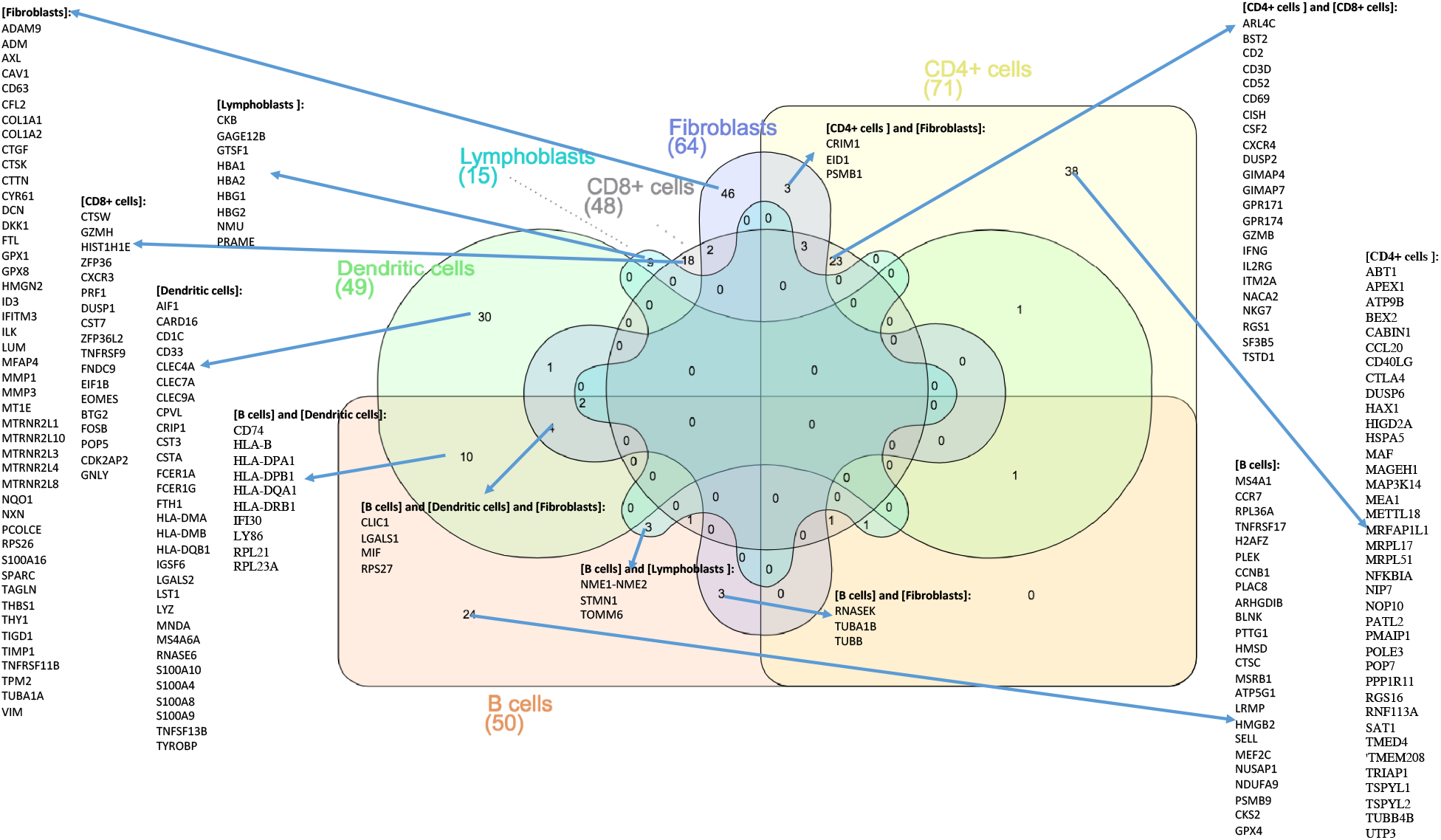
Venn diagram representation of genes with p-value of less than 0.01 for each of the six cell-types by MICTI. *The Venn diagram was created using InteractiVenn* (Heberle, Meirelles, da Silva, Telles, & Minghim, 2015).

23 genes were found to be cell-type specific genes for both CD4+ and CD8+ memory cells (Fig. 5). They were enriched for Th1 and Th2 cell differentiation, JAK-STAT signalling pathway, T cell receptor signalling pathway, Th17 cell differentiation and Natural killer cell mediated cytotoxicity (*supplement file 7*) KEGG pathways. It is known that the Th1, Th2 and Th17 cells are sub-populations of the CD4+ helper T cells. CD8+ (cytotoxic) T cells are also involved in cell killing by secreting enzymes into the infected cell. There were 38 genes identified as exclusively CD4+ cell-type specific, while there were 18 genes that were identified as exclusively CD8+ cell markers. The exclusive CD4+ cell-type marker genes were enriched for T cell receptor signalling pathway (*supplement file 7*).

There were 49 dendritic cell-type specific genes as it is shown in Fig. 5. The gene ontology terms associated with these genes were antigen processing and presentation, immune response, leukocyte activation, cell activation, leukocyte activation involved in immune response, cell activation involved in immune response, MHC class II protein complex assembly and T cell activation (*supplement file 7*) among others. (Banchereau et al., 2000) has shown that these ontology terms are of known dendritic cell’s molecular functionality.

Fig. 5 also shows 64 genes that were specific to fibroblast cells. These genes were enriched with biological terms of extracellular structure organization, extracellular matrix organization, response to wounding and wound healing (*supplement file 7*). Fibroblast cells are found in most tissues of the body and these ontology terms explain the typical fibroblastic cellular function as it was detailed in (McAnulty, 2007). The lymphoblast cells that we used in our analysis were hematologic cancerous cells with Chronic myeloid leukemia (CML). Our algorithm found 15 genes that were specific to this group of cells, out of which, nine of them were exclusive lymphoblast specific genes. These genes were enriched for gas transport, drug transport and oxygen transport biological ontology terms (*supplementary file 7*)

Finally, we compared ***MICTI*** in terms of execution time with the other pairwise statistical testing methods for differential expression analysis using MAST (Finak et al., 2015), DESeq2 (Love, Huber, & Anders, 2014), ROTS (Suomi et al., 2017) and BPSC (Vu et al., 2016) pipelines.

Fig. 6 indicates the time it takes for pairwise comparisons of the one group of cell-type against the other cell-types for each of underlined statistical testing methods. DESeq2 is expensive in terms of execution time taking about nine hours per each of the statistical testing while BPSC is taking about two hours per statistical testing. ROTS, a dataset independent statistical testing pipeline, is relatively fast taking about 12 minutes per statistical testing. MAST took about 3 minutes per pairwise statistical test for differential expression. ***MICTI***, which is a feature extraction method for marker gene identification, only takes about less than two seconds for all cell-type specific gene identification for each of cell-types. The comparison is performed on CSC EC2 instance with Ubuntu 16.04.5 OS, 29.3GB of RAM and 8 virtual CPUs with the capacity of 2GHz for each.

**Figure 6.**
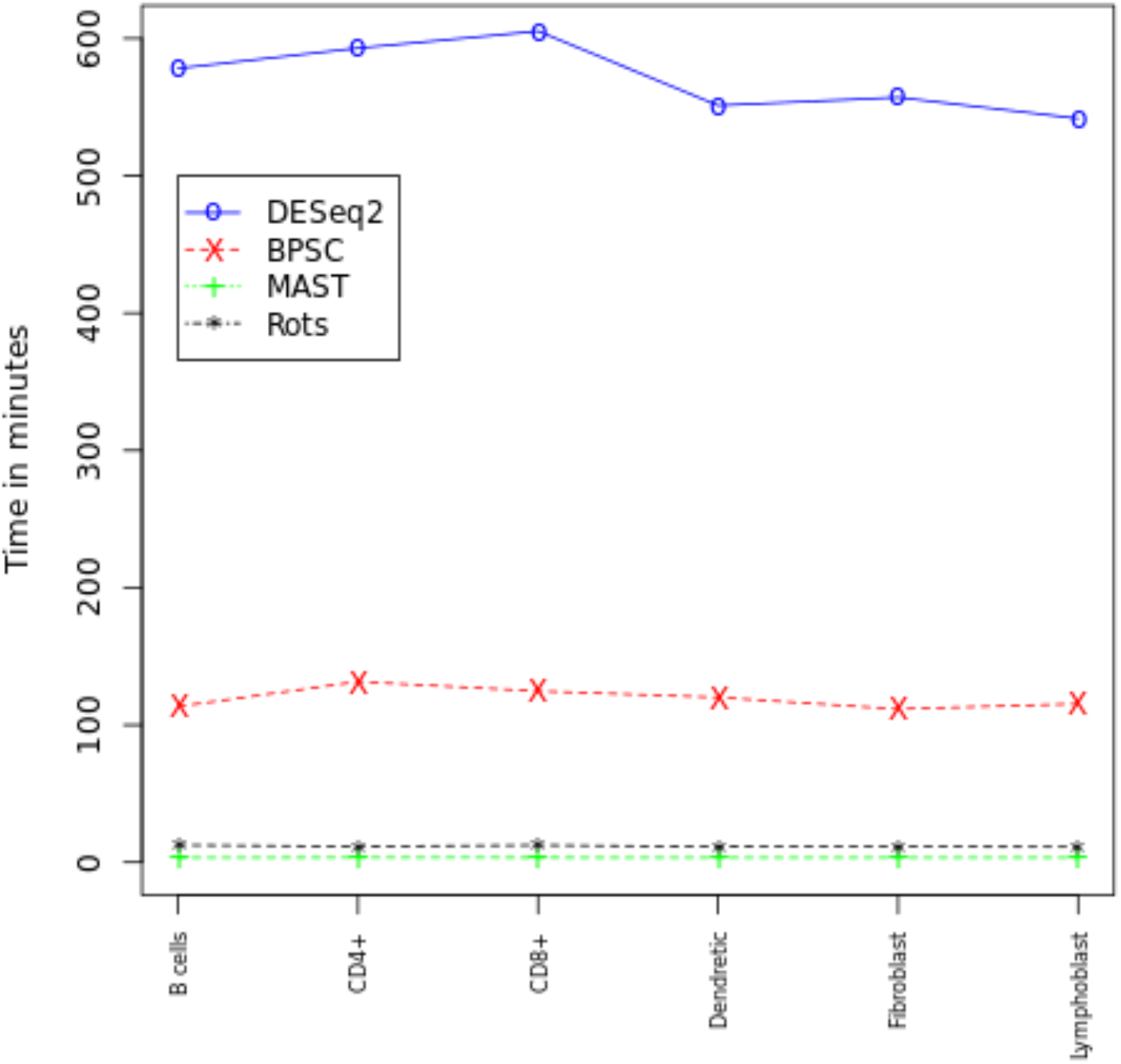
Comparison of different statistical differential expression pipelines for single-cell data in terms of execution time for cell-type specific marker gene identification. The x-axis is the pairwise testing of each cell-type verses the other cell-types. The y-axis represents the execution time for each of pairwise statistical testing in minutes.

## Discussion

We proposed ***MICTI*** as a feature extraction method for key marker gene identification in the heterogonous single-cell RNA-seq data. The nature of single-cell RNA-seq data is the underlying cell heterogeneity. Clustering of cells by their cell-types using unsupervised machine learning methods usually results in unequal numbers of cells across the clusters. This has consequences for differential gene expression analysis as the statistical power and type I error is highly dependent on the number of samples and the variance of gene expression values in the compared clusters (Rusticus & Lovato, 2014). The statistical power decreases as the sample sizes become unequal. If both the sample sizes and variances are unequal, this can result in dramatic difference in power and the type I error rate gets inflated or reduced (Rusticus & Lovato, 2014).

In addition, pairwise statistical testing, using for example *BPSC and DEseq2(*Fig. 6), for each of the cell-types against the other cell-types in the population take much execution time. Moreover, there is no a gold standard method for differential expression analysis in the context of single-cell RNA-seq data (Dal Molin, Baruzzo, & Di Camillo, 2017). Our feature extraction based method, which differs from pairwise statistical testing methods in using cell-type specific mean feature-encoded expression values for marker gene significance *p*-value calculation, can efficiently identify key cell-type specific genes from heterogeneous cell population expression data. It also avoids gene filtering based-on expression variance, a common procedure before proceeding to the downstream clustering and differential analysis that might result in loss of important information, in most of single-cell RNA-seq analysis pipelines. It also has significantly less execution time when it is compared with the statistical testing methods(Fig. 6).

***MICTI*** does not depend on balanced numbers of cells in each of clusters. In the case studies 1, 2 and 3, the number of cells varied in each of the heterogeneous cell categories (Fig. 3a and Fig. 4a), but ***MICTI*** identified key marker genes in each cluster or cell-types. We can also observe that the ***MICTI*** identified cell-type marker genes for functionally related cell-types, such as B-cells and Dendritic cells or CD4+ and CD8+ memory cells, share communality (Fig. 4b).

Hence, we suggest ***MICTI*** as alternative or supplementary method for users working with the heterogeneous single-cell RNA-seq dataset. Finally, there are massive single-cell RNA-seq data being produced as the technology advances. The discovery of the new cell-types and their underlying biological significance can be unfolded by incorporating machine learning. As feature extraction is a key step in machine learning, ***MICTI*** can be used for the large scale machine learning task for cell-type classification.

## Conclusions

***MICTI***, feature extraction method for marker gene identification for single-cell RNA-seq data, can be used as an alternative to statistical differential expression analysis for marker gene identification. It not only avoids repeated statistical testing for marker gene identification in the heterogeneous cell population, but it also keeps gene expression information otherwise lost in filtering genes based on their variance across the cells as it is common practice in most single-cell RNA-seq pipelines.

## Methods

***MICTI*** is a feature extraction-based method for identification of cell-type specific marker genes in single-cell RNA-seq data. We describe the typical workflow of ***MICTI*** in detail. Following grouping of cells into their respective cell-type categories, in our case we did it manually by extracting cell metadata information, the next step was to investigate the marker genes that gave rise to the identity of the cell in the heterogeneous cell population. This was done by looking at the gene expression level of a given gene in each cell weighted by the scarcity of expression factor for the given gene across the cell-types. The scarcity of expression factor for a given gene was calculated by dividing the total number of cells by the number of cells that express the given gene in the logarithmic scale. We considered a gene as “expressed” if the normalized UMI count or TPM value was greater than the user-defined threshold value as the binary Yes/No decision. We call this step as “feature-encoding” for subsequent feature extraction as (equations 1-5). Fig. 1 shows the workflow of this feature extraction approach for identifying cell-type specific genes or markers.

### Feature encoding

We transformed the normalized expression value of each gene in every cell by encoding how scarcely the given gene was expressed across all cells. The gene expression scarcity factor was calculated by dividing the total number of cells by the number of cells that expressed the given gene in the logarithmic scale. The threshold expression value that decides whether a gene is expressed or not is given by the user. By default, we assigned it to be 0, i.e., genes with expression value of 0 were considered as non-expressed genes while the ones that had greater than 0 values were considered as expressed in their respective cells. The transformed expression values were the product of the normalized expression value and the gene expression scarcity factor for each gene in every cell. Thus, we encoded the cell-type specificity for the given gene across all the cells. A more detailed description for ***MICTI*** workflow is given below.

Before applying the feature encoding on the gene expression values, the expression values of each gene should be normalized. For the non-UMI data, the TPM normalization is adopted as it allow cross cell gene expression comparison adjusted by gene length. Considering the UMI-based data, the raw UMI count *UMI_ij_* of gene *i* within each cell *j* can be normalized by:

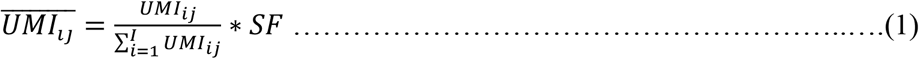

where *SF* is a scaling factor to scale the otherwise very small relative expression values (*SF, recommended 10^6^*) for ease of computation. The normalized expression value 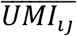 of each gene was then transformed to a feature-encoded expression value *fe_ij_* by multiplying it with the scarcity of expression factor 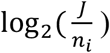:

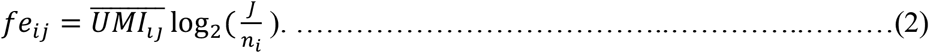

The scarcity of expression factor is the logarithmic ratio of the total number of cells *J* to the number of cells *n_i_* that express the particular gene *i* across the cells with a user-defined threshold value *θ*:

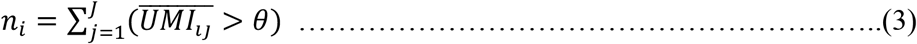

A suitable threshold value depends on the library size and the dataset. We used the threshold expression value of 0.

### Marker gene identification

We calculated the cluster level feature-encoded gene expression values *F_i,k_* for each gene *i* and cluster *k* by calculating the mean feature-encoded expression value of the gene in the cluster divided by the cluster variance in the logarithmic scale:

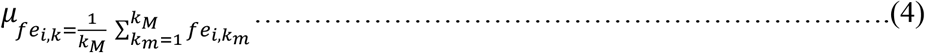

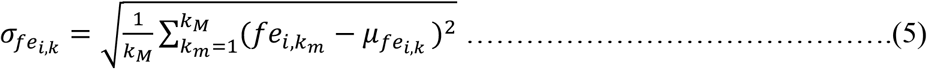

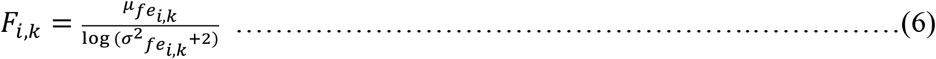

where *k_M_* is the total number of cells in cluster *k, μ_fe_i,k__* is the mean feature-encoded expression values for gene *i* among cells in cluster *k, σ_fe_i,k__* is the standard deviation of the feature-encoded expression values for gene *i* within the cluster *k. F_i,k_* is the normalized cluster mean feature-encoded value for the given gene *i* in a cluster *k*. The normalization of cluster mean feature-encoded values with the cluster variance allow the identification of markers that are common for most of the cluster members avoiding the bias towards the high mean feature-encoded expression value from the outlier cells. Finally, the *Z*-score for each gene in the given cluster was calculated:

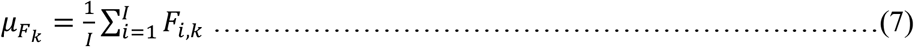

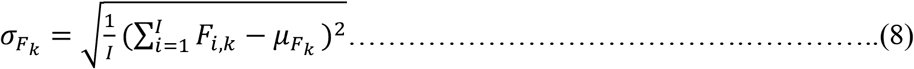

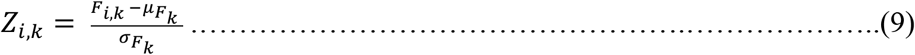

where *μ_F_k__* is the average normalized feature-encoded value of all genes *I* in a given cluster *k, σ_F_k__* is standard deviation of the normalized feature-encoded expression values for the given cluster *k*. The positive extreme Z-score valued genes were cluster-specific genes, whereas the negative extreme Z-score valued genes were the expressed genes across all clusters. In order to identify the statistically significant cluster specific marker genes, we calculated the Bonferroni corrected *p-values* for each of the clusters. We used 0.01 as a cut-off p-value, where the significant genes with *p-values* of less than 0.01 and *Z*-score of greater than 0 were cluster markers.

### Gene list enrichment analysis

In order to validate cell-type specific marker genes, we used gene over-representation analysis for pathways and gene ontology (GO) terms. For the over representation analysis, we used *gprofiler(version 1.1.0)*(Reimand, Arak, & Vilo, 2011), a python package for analysing the gene over-representation. *gprofiler* uses set counts and sizes (SCS) method to estimate threshold *p-values* from complex and structured functional profiling data such as GO, pathways and TFBS where the statistical significance is determined from set intersections in 2×2 contingency table (Reimand, Kull, Peterson, Hansen, & Vilo, 2007).

### Dataset and data preparation

#### Case study 1

We used SymSim (Zhang et al., 2019), an R software package, to generate simulated single-cell RNA-seq data for five distinct cell populations with the tree structure represented in Fig. 2a. (Zhang et al., 2019) implemented a classical promoter kinetic model with kinetic parameters for promoter on rate, *k_on_*, promoter off rate, *k_off_* and RNA synthesis rate, s to generate the true transcript counts accounting for the extrinsic, intrinsic and technical variations.

The paraments used for the generation of these UMI based transcript count dataset were the number of genes (ngenes=5000), the number of cells (ncells=200), minimum population size for the given groups (min_popsize=30), the smallest population size (i_minpop=2), the value from which the extrinsic variability factor(evf) mean is generated from (evf_center=1), the number of evf for each kinetic parameter (nevf=10), the population structure of the cells (evf_type=“discrete”), the number of differential evfs between populations for one kinetic parameter (n_de_evf=9), determining the kinetic parameters with differential evfs (vary=“s”), controls parameter for heterogeneity of cells in each of the cell population (Sigma=0.5), the parameter that controls difference between genes (gene_effects_sd=1), the probability of non-zero values of genes in the gene effect vectors (gene_effect_prob=0.3), parameter that adjusts for bimodality of gene expression for controlling intrinsic variation (bimod=0), random seed to reproduce the results (randseed=0), the experimental dataset used to estimate the kinetic parameters “*k_on_*”, “*k_off_*” or “*s*” (param_realdata = “zeisel.imputed”), parameters that determines the return format of simulated data weather it is in the form of summarized experiment or list of elements format (SE=F). We used within cell library size normalization for each of the cells in the dataset.

#### Case study 2

We used TPM normalized, publicly available human pancreatic islet single-cell RNA-seq data with the GEO accession number of GSE86469.

#### Case study 3

All single cell gene expression datasets used in this study were publicly available under GEO accessions GSE75748, GSE81861, GSE44618, GSE81861, GSE96562, GSE96563, GSE96568, GSE85527, GSE85530, GSE96564 and GSE89232. We used the pre-processed count matrices which were generously provided by the corresponding authors. To be able to compare expression values across different experiments and platforms, we normalized the expression values of each gene with the library size of each cell, as each of the cells had different library size across the experiments. We chose to use TPM normalization with a requirement that the total read count for all genes per cell sum to 10^6^. This allowed us to perform pairwise comparisons of gene expression levels across all cell-types. Then, we kept genes that had greater than zero TPM normalized value in at least one of the cells in the 11 experimental data mentioned above. Finally, we proceeded with the TPM normalized expression values of 13880 genes in 1153 cells.

### Software availability

Software used to generate all analyses in this manuscript is publicly available as a PYPI python package (https://pypi.org/project/MICTI/) and included here as a supplementary Software in GitHub.

## List of abbreviations

MICTI: *M*arker gene Identification for *Cell-type I*dentity
DE: Differential Expression
MAST: Model-based Analysis of Single-cell Transcriptomics.
ROTS: The Reproducibility-Optimized Test Statistic
BPSC: Beta-Poisson model for Single-Cell RNA-seq data analyses
EC2: Elastic cloud compute
DGE: Differential Gene Expression
TPM: Transcript Per Millions of mapped read
RPKM: Read Per Kilobase per Millions of mapped read
GEO: Gene Expression Omnibus
UMI: Unique Molecular Identifier
SC3: Single-Cell Consensus Clustering
LDA: Latent Dirichlet Allocation
PCA: Principal Component Analysis
ICA: Independent Component Analysis
TF-IDF: Term Frequency Inverse Document Frequency
NMF: Negative Matrix Factorization
t-SNE: t-Distributed Stochastic Neighbor Embedding
MST: Minimum Spanning Tree
TSCAN: Tools for Single-Cell Analysis
NB: Negative Binomial
FACs: Fluorescence-Activated Cell Sorter
PCR: Polymerase Chain Reaction
cDNA: complementary DNA
scRNA-seq: Single-Cell RNA Sequencing

## Declarations

### Ethics approval and consent to participate

Not applicable.

### Consent for publication

Not applicable.

### Authors’ contributions

## Acknowledgements

The authors wish to acknowledge CSC – IT Center for Science, Finland, for computational resources.

